# The permeabilized SecY protein-translocation channel can serve as a nonspecific sugar transporter

**DOI:** 10.1101/378786

**Authors:** Sen Mei, Chong Xie, Hao Mi, Chuang Xue, Qiang Guo, Guang-Qing Du, Guo-Bei Li, Cai-Xia Li, Ya-Nan Qu, Ming-Hao Xiong, Yang Jiang, Tian-Wei Tan, Shang-Tian Yang, Li-Hai Fan

**Author notes:** Sen Mei and Chong Xie contributed equally to this work. Corresponding author: Li-Hai Fan; Address: North Third Ring Road 15, Chaoyang District, Beijing, People’s Republic of China, 100029.

## Abstract

As the initial step in carbohydrate catabolism in cells, the substrate-specific transporters via active transport and facilitated diffusion play a decisive role in passage of sugars through the plasma membrane into the cytoplasm. The SecY complex (SecYEG) in bacteria forms a membrane channel responsible for protein translocation. This work demonstrates that weakening the sealability of the SecY channel allowed free diffusion of sugars, including glucose, fructose, mannose, xylose, arabinose, and lactose, into the engineered cells, facilitating its rapid growth on a wide spectrum of monosaccharides and bypassing/reducing stereospecificity, transport saturation, competitive inhibition, and carbon catabolite repression (CCR), which are usually encountered with the specific sugar transporters. The SecY channel is structurally conserved in prokaryotes, thus it may be engineered to serve as a unique and universal transporter for bacteria to passage sugars as demonstrated in *Escherichia coli* and *Clostridium acetobutylicum*.

## Introduction

Synthetic biology aims to design and build biological systems to reprogram organisms or even create artificial life (Smanski *et al*, 2016; Ausländer *et al*, 2017). Sugars are the major carbon and energy sources for living cells, thus transport of sugars across the plasma membrane into the cytoplasm is an essential cell physiological function, which relies on the sugar-specific transport systems involving active and passive mechanisms (Cirillo, 1961; Reinhold & Kaplan, 1984; Chen *et al*, 2015). Active transport requires metabolic energy to passage molecules across the plasma membrane in the direction against the concentration gradient, whereas passive transport moves molecules down its concentration without needing an energy input, which can be further classified as facilitated diffusion and free diffusion, respectively. Active transport and facilitated diffusion are dependent on the specific molecular binding between cargo and transporter, thereby exhibiting stereospecificity, transport saturation, and competitive inhibition (Cirillo, 1961). Moreover, numerous sugars are taken up with concomitant phosphorylation via the phosphoenolpyruvate (PEP): carbohydrate phosphotransferase system (PTS), which may cause carbon catabolite repression (CCR), leading to a diauxic cell growth (Stülke & Hillen, 1999). Fig 1A illustrates the main transporters of glucose, fructose, mannose, xylose, arabinose, and lactose in *Escherichia coli*, a Gram-negative, facultatively anaerobic bacterium. In contrast, free diffusion is a non-mediated transport with a linear rate in proportion to the concentration difference. However, it is believed that only small, non-polar molecules, such as carbon dioxide and oxygen, can freely diffuse through the plasma membrane (Zhao *et al*, 2011). Therefore, although many sugars are transported passively, no sugars are known to enter cells via a free diffusion manner (Romano, 1986).

**Figure 1.**
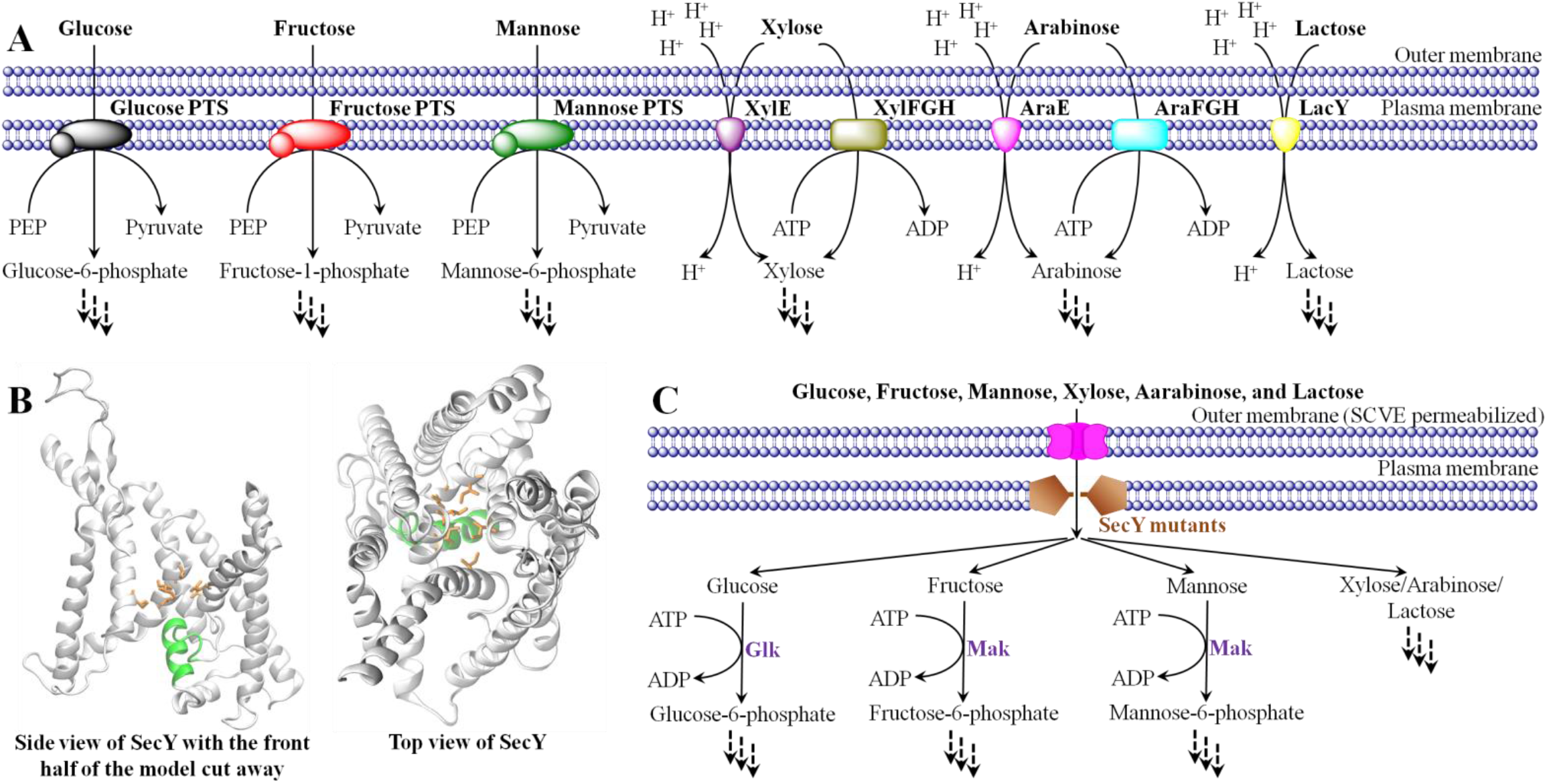
Use of SecY mutants for sugar transport across the plasma membrane of *E. coli*. **A.** The main uptake routes of glucose, fructose, mannose, xylose, arabinose, and lactose in wild-type *E. coli*. **B.** The 3D structure of the wild-type SecY channel obtained via homology modeling (Modeller v9.18) using PDBID 3J46 as a template. The plug domain is shown in green, and the pore ring is orange. The pore-ring residues of SecY: Ile 82, Ile 86, Ile 187, Ile 191, Ile 278, Ile 408. **C.** Nonspecific transport of sugars via SecY mutants.

The protein translocation channel is a structurally conserved protein-conducting system that allows polypeptides to be transferred across or integrated into cell membranes (Osborne *et al*, 2005; Rapoport, 2007; Mandon *et al*, 2009). In bacteria, this channel is formed by the SecY complex (SecYEG) and is located in the plasma membrane (Brundage *et al*, 1990; Akimaru *et al*, 1991; Breyton *et al*, 2002). The X-ray crystal structure revealed that the SecY channel has an hourglass shape with a narrow central constriction consisting of six hydrophobic amino acid residues termed the pore ring, and the exofacial side of the channel is occluded by a short α-helical segment called the plug domain (Van den Berg *et al*, 2004) (Fig 1B). The SecY plug domain serves as a physical barrier that ensures the channel is sealed in its resting state (Li *et al*, 2007). During protein translocation, the plug is displaced away from the center of the channel toward the open position by the incoming polypeptide (Tam *et al*, 2005; Li *et al*, 2016), and the pore ring forms a ‘gasket-like’ seal around the polypeptide chain (Park & Rapoport, 2011), preventing the permeation of other molecules.

Given the widespread interests in sugar transporters in basic biology research and biotechnology application (Galazka *et al*, 2010; Luo *et al*, 2014), as well as the challenges surrounding their broad substrate profiles (Kornberg *et al*, 2000), these findings prompted us to hypothesize that whether the SecY channel can be designed as a free diffusion passageway for efficient transport of carbohydrates, thus developing an entirely novel paradigm for sugar passage and addressing the problems encountered with the specific transporters. In the present work, we engineered the SecY channel by plug deletion and pore ring mutation, and demonstrated that nonspecific transport of mono- and di-saccharides inward across the plasma membrane could be achieved without requiring cells to possess a large number of specific sugar transporters for survival by using the SecY channel mutants in combination with the SARS (severe acute respiratory syndrome) coronavirus envelope protein (SCVE) and the appropriate sugar kinase (Fig 1C), thereby overcoming a bottleneck in engineering bacteria for growth on various sugars, a desirable attribute for biotechnology applications.

## Results and Discussion

### Functional transport of the PTS-dependent hexoses via SecY (ΔP) Channel

When we expressed the SecY with deletion of the plug (SecY (ΔP)) in an *E. coli* (ΔPtsG) lacking the glucose PTS (Plumbridge, 1998) (Fig 2A and B), the cell growth defect was not observed (Fig 2D and E). In contrast, the limited growth on glucose caused by insufficient import of the sole carbon source was partially rescued. We reasoned that the SecY (ΔP) channel might allow glucose to diffuse through the plasma membrane due to its small *V*_A_ (≈ 0.178 m^3^/kmol) and large diffusion coefficient (*D*_AB_ ≈ 1.009×10^−9^ m^2^/s) in water (Appendix Table S1), which was confirmed by a [^14^C]-labelled glucose transport assay (Fig 2C). With the purpose of further enhancing the glucose uptake, SCVE was co-expressed with the SecY (ΔP) channel (Fig 2A and B). The SCVE has the ability to permeabilize the outer membrane of *E. coli* by formation of the transmembrane pores as porins (Liao *et al*, 2004). The reduced permeation barrier of the outer membrane might increase the glucose gradient across the SecY (ΔP) channel, thus accelerating the glucose diffusion into the cytoplasm (Fig 2C) and resulting in a marked improvement of the growth of *E. coli* (ΔPtsG) on glucose (Fig 2D and E) as well as the wild-type strain (Fig EV1). Moreover, the glucose PTS catalyzes the transport with concomitant phosphorylation of glucose to glucose-6-phosphate. Therefore, once glucose was inside the mutant *E. coli*, phosphorylation might become the activity limiting glycolytic flux (Hernández-Montalvo *et al*, 2003), which could explain the total recovery of the growth of *E. coli* (ΔPtsG) on glucose by the over-expression of an *E. coli* glucokinase (Glk) (Fukuda *et al*, 1983) together with SecY (ΔP) and SCVE (Fig 2D and E). We compared the Monod growth kinetic parameters of the *E. coli* (ΔPtsG) co-expressing SecY (ΔP), SCVE and Glk with those of the wild-type strain (Fig 2F). Both strains had a maximum specific growth rate (*μ*_max_) of 0.347 h^−1^, but the half-saturation constant (*K_S_*) of the mutant strain (≈ 4.803 mM) was 1.49-fold that of the wild-type *E. coli* (≈ 3.234 mM).

**Figure 2.**
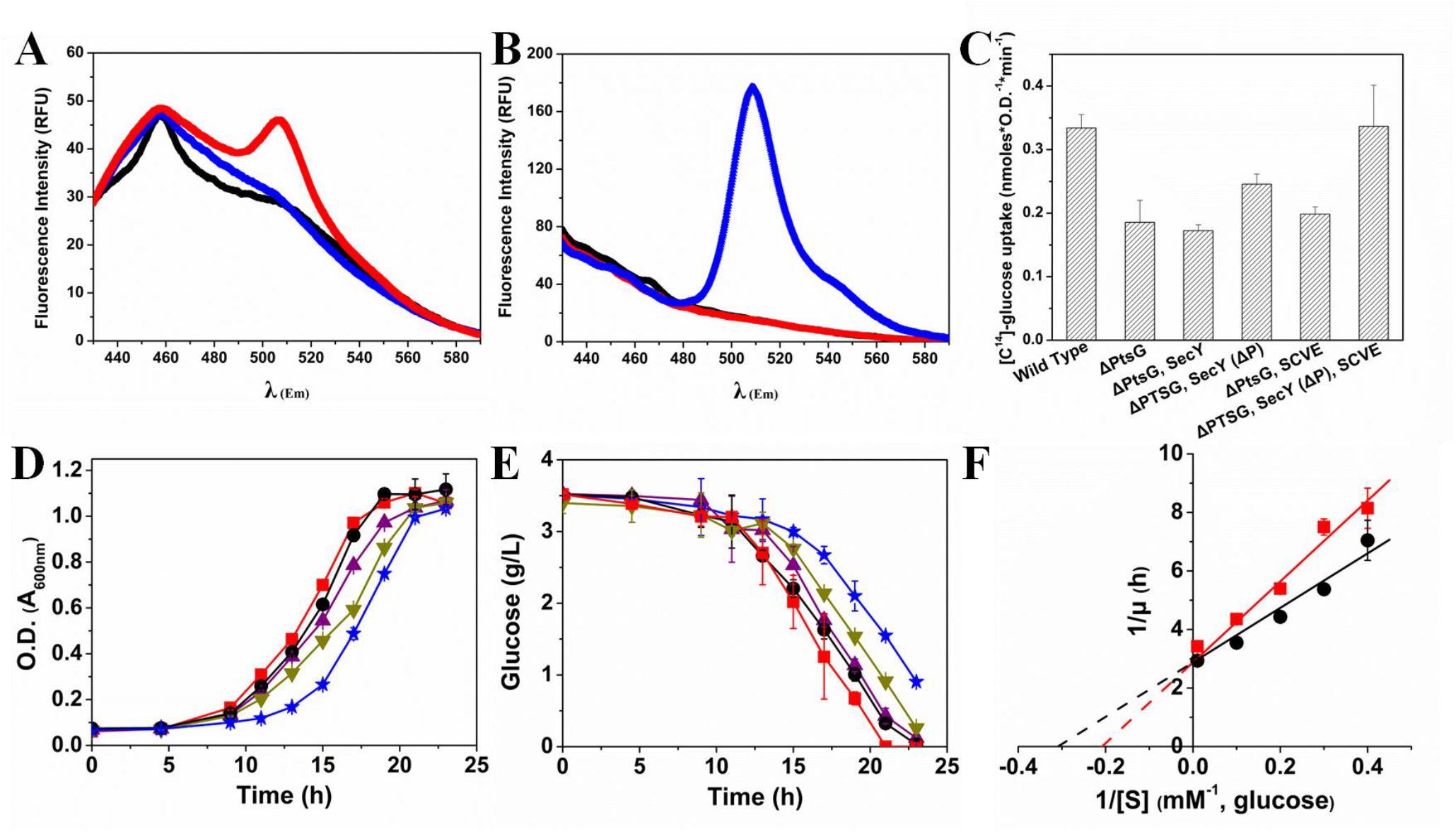
The SecY (ΔP) channel allows glucose to enter the *E. coli* cells via free diffusion. **A, B.** Expression and localization of SecY (ΔP) and SCVE in *E. coli* (ΔPtsG). (A) Emission scanning of the plasma-membrane fraction, (B) Emission scanning of the outer-membrane fraction. *E. coli* (ΔPtsG) (black), *E. coli* (ΔPtsG) expressing the GFP-fused SecY (ΔP) (red), *E. coli* (ΔPtsG) co-expressing SecY (ΔP) and the GFP-fused SCVE (blue), *λ*_Ex_ = 395 nm. **C.** Uptake of [C^14^]-glucose by *E. coli*. SecY means the cell over-expressing the wild-type SecY channel. Error bars, s.d., n=3. **D, E, F.** The growth kinetics of *E. coli* strains in M9 medium with glucose. Wild-type *E. coli* (black circle), *E. coli* (ΔPtsG) (blue star), *E. coli* (ΔPtsG) with SecY (ΔP) (dark yellow triangle), *E. coli* (ΔPtsG) with SecY (ΔP) and SCVE (purple triangle), *E. coli* (ΔPtsG) with SecY (ΔP), SCVE and Glk (red square). It should be noted that PtsG deletion reduced the cell growth rate instead of preventing cell growth, since the glucose transport pathways could be shifted to ManXYZ, EIIB’^Fru^/EIIBC^Fru^ and/or EIIC^*bgl*F3^. Error bars, s.d., n=3.

Then, we selected fructose and mannose as two other models of the PTS-dependent hexoses. *E. coli* with damaged fructose PTS (ΔFruA) (Ferenci & Kornberg, 1974) or mannose PTS (ΔManXYZ) (Erni & Zanolari, 1985) did not grow on fructose (Fig 3A and B) or mannose (Fig 3D and E), respectively. On the other hand, SecY (ΔP) and SCVE were found to help both mutant strains to recover cell growth, especially for the one grown on mannose. By co-expression of an *E. coli* fructo(manno)kinase (Mak) which catalyzes the phosphorylation of fructose or mannose in the cytoplasm (Sproul *et al*, 2001), the strain lacking FruA grew on fructose at a similar rate to that of the wild-type *E. coli* (Fig 3A and B), and the growth of the strain without ManXYZ on mannose was even faster than the control (Fig 3D and E), suggesting that the fructose or mannose imported by the SecY (ΔP) channel already satisfied the cell demand. It should be noted that the fructose taken up via the fructose PTS is concomitantly phosphorylated to fructose-1-phosphate before further phosphorylation to fructose-1,6-bisphosphate (Kornberg, 2001) (Fig 1A), but in our route, the phosphorylated fructose first appears inside the cell as fructose-6-phosphate (Fig 1C), which is in agreement with the route using the transporter of PtsG-F (Kornberg *et al*, 2000). The data of growth kinetics in Fig 3C and F show that co-expression of SecY (ΔP), SCVE and Mak enabled the mutant *E. coli* with almost the same *μ*_max_ values as those of the wild-type strain, which were 0.248 h^−1^ on fructose and 0.398 h^−1^ on mannose. Also, as in the case of glucose, the SecY (ΔP) channel exhibited a higher half-saturation constant for fructose (*K_S_* ≈ 1.579 mM) and mannose (*K_S_* ≈ 4.719 mM) than the fructose PTS (*K_S_* ≈ 0.843 mM) and the mannose PTS (*K_S_* ≈ 3.181 mM), respectively.

**Figure 3.**
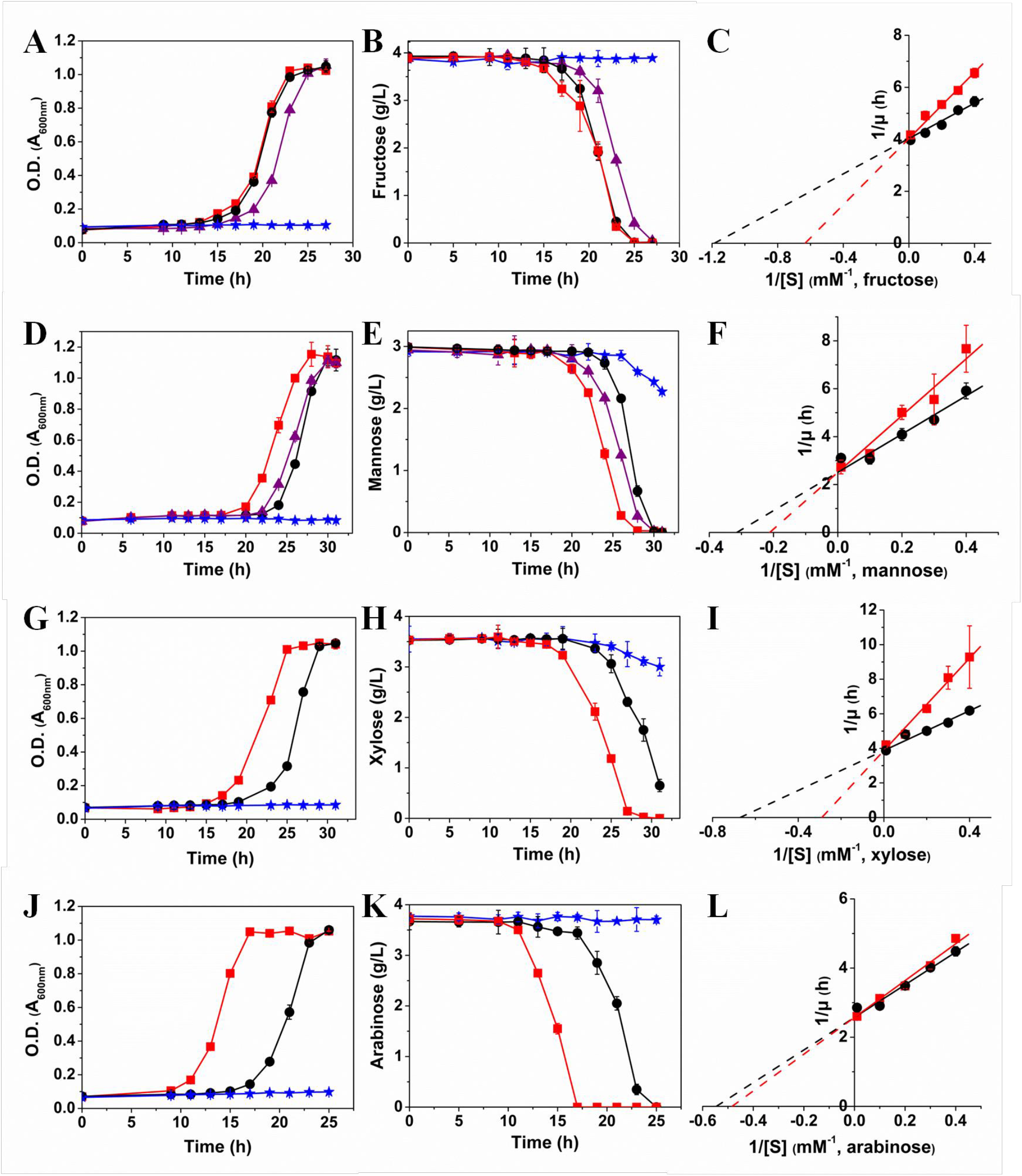
Comparison of the growth of the wild-type and the engineered *E. coli* in M9 medium with various sugars. **A, B, C.** The growth kinetics of *E. coli* strains on fructose. Wild-type *E. coli* (black circle), *E. coli* (ΔFruA) (blue star), *E. coli* (ΔFruA) with SecY (ΔP) and SCVE (purple triangle), *E. coli* (ΔFruA) with SecY (ΔP), SCVE and Mak (red square). Error bars, s.d., n = 3. **D, E, F.** The growth kinetics of *E. coli* strains on mannose. Wild-type *E. coli* (black circle), *E. coli* (ΔManXYZ) (blue star), *E. coli* (ΔManXYZ) with SecY (ΔP) and SCVE (purple triangle), *E. coli* (ΔManXYZ) with SecY (ΔP), SCVE and Mak (red square). Error bars, s.d., n = 3. **G, H, I.** The growth kinetics of *E. coli* strains on xylose. Wild-type *E. coli* (black circle), *E. coli* (ΔXylFGH) (blue star), *E. coli* (ΔXylFGH) with SecY (ΔP) and SCVE (red square). Here, damage of XylFGH was found to prevent cell growth, so XylE was not co-deleted. Error bars, s.d., n = 3. **J, K, L.** The growth kinetics of *E. coli* strains on arabinose. Wild-type *E. coli* (black circle), *E. coli* (ΔAraFGH, ΔAraE) (blue star), *E. coli* (ΔAraFGH, ΔAraE) with SecY (ΔP) and SCVE (red square). Error bars, s.d., n = 3.

### Functional transport of the non-PTS-dependent pentoses via SecY (ΔP) channel

Next, we tested xylose and arabinose, which are non-PTS-dependent pentoses. The main transporters for xylose (XylFGH) (Sumiya *et al*, 1995) and arabinose (AraFGH, AraE) (Schleif, 2000) in *E. coli* were deleted independently, thus the obtained strains either did not grow on xylose (Fig 3G and H) or arabinose (Fig 3J and K) as the sole carbon source. As expected, co-expression of SecY (ΔP) and SCVE fully supported the growth of the mutants. The lag phases of both mutant strains were obviously shortened when compared with the wild-type *E. coli* (Fig 3G and J), which was also found in the case of mannose (Fig 3D). Interestingly, previous work reported that a mutant glucose facilitator protein (2-RD5) from *Zymomonas mobilis* allowed rapid transport of xylose in *E. coli*, but in contrast, 2-RD5 caused an extensive growth delay (Ren *et al*, 2009). A prolonged lag phase (≈ 25 h) on arabinose was also observed when we re-expressed AraFGH together with SCVE in *E. coli* (ΔAraFGH, ΔAraE) (Fig EV2), even though AraFGH is regarded as an efficient uptake system for arabinose (Luo *et al*, 2014). The use of the SecY (ΔP) channel did not seem to facilitate the initiation of the arabinose-inducible promoter *P_BAD_* (Fig EV3), which controls the synthesis of the key enzymes (AraBAD) involved in the pathway of arabinose metabolism in *E. coli* (Schleif, 2000). Therefore, we suspect that the SecY (ΔP) channel might allow *E. coli* to import not only sugars but also other small nutritional molecules, for which the cells adapted to growth conditions more quickly than the wild-type strain. The mutant strain expressing SecY (ΔP) with SCVE had *μ*_max_ ≈ 0.258 h^−1^ on xylose (Fig 3I) and ≈ 0.388 h^−1^ on arabinose (Fig 3L), which were almost identical to those of the wild-type *E. coli*. The *K_S_* values for the mutant strains were 3.449 mM and 2.066 mM on xylose and on arabinose, respectively, while for the wild-type *E. coli*, the values were 1.489 mM and 1.831 mM.

### Functional transport of disaccharide via SecY (ΔP, IIIGIG) channel

After that, we investigated the transport of lactose as a model disaccharide via the SecY (ΔP) channel. The LacY transporter (Abramson *et al*, 2003) was deleted from *E. coli*, thus the obtained strain could not grow on lactose as the sole carbon source (Fig 4A). However, only a slight growth was observed by co-expression of SecY (ΔP) and SCVE, probably because the larger molecular volume of lactose (*V_A_* ≈ 0.340 m^3^/kmol) leads to a lower diffusion coefficient (*D*_AB_ ≈ 0.684×10^−9^ m^2^/s) than monosaccharides (*V*_A_ ≤ 0.178 m^3^/kmol, *D*_AB_ ≥ 1.009×10^−9^ m^2^/s) (Appendix Table S1), resulting in a reduced sugar flux through the pore ring of SecY (ΔP). We thus replaced the isoleucines (molecular weight *Mr* ≈ 131.18 g/mol) in the pore ring with glycines (*Mr* ≈ 75.07 g/mol) to amplify the channel open size (Fig 4B). As illustrated in Fig 4C, the glycine substitutions at Ile 191 and Ile 408 (SecY (ΔP, IIIGIG)) provoked a 1.27-fold increase in permeation of 2-deoxy-2-[(7-nitro-2,1,3-benzoxadiazol-4-yl)amino]-D-glucose (2-NBDG) which has almost the same *V*_A_ and *D*_AB_ as lactose (Appendix Table S1). By co-expression of SecY (ΔP, IIIGIG) and SCVE, *E. coli* (ΔLacY) exhibited a moderate growth on lactose 50 h after cell inoculation (Fig 4A).

**Figure 4.**
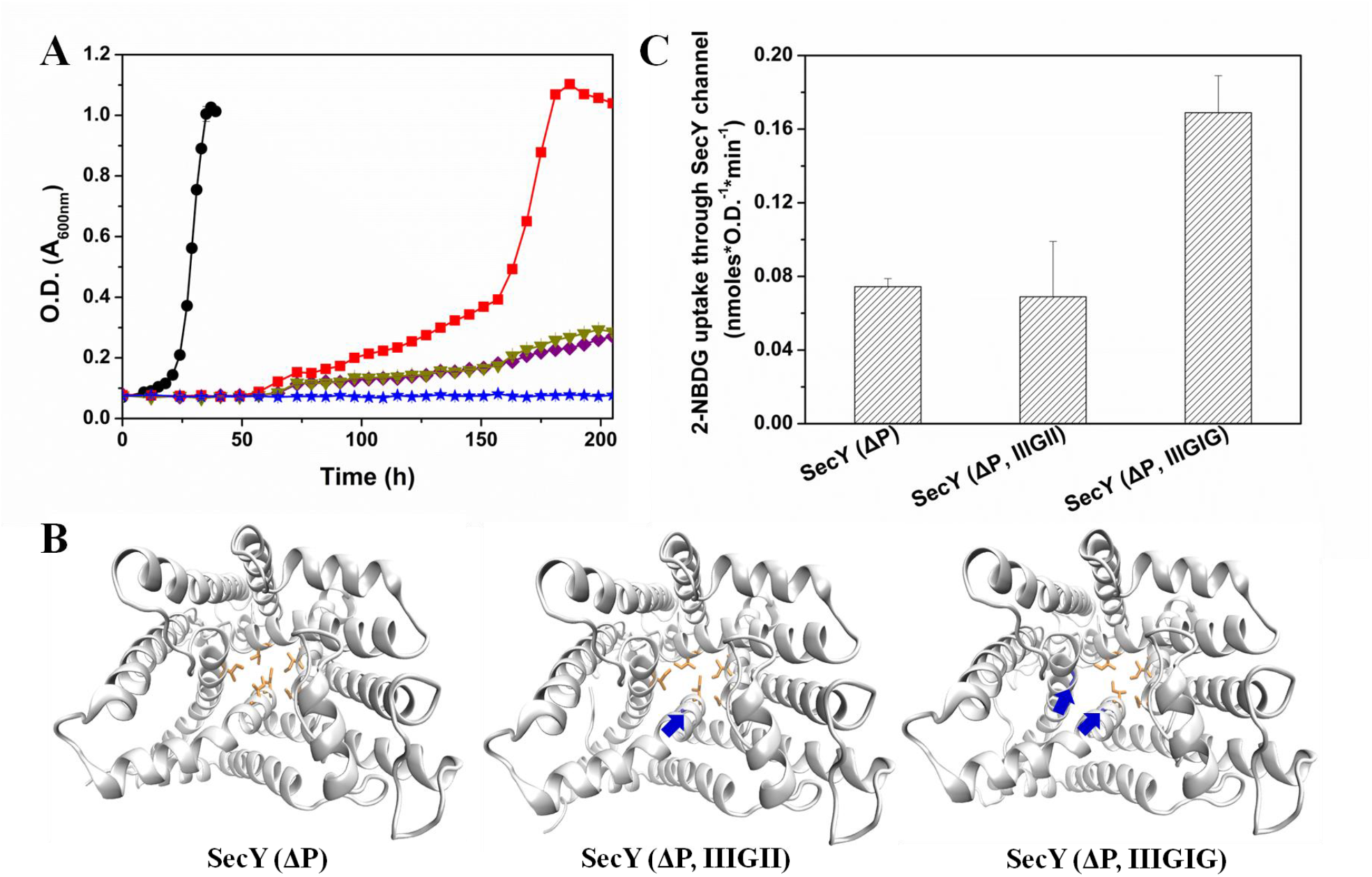
Use of SecY(ΔP, IIIGIG) for lactose transport across the plasma membrane of *E, coli*. **A.** Cell growth in M9 medium containing 0.4% (w/v) lactose. Wild-type *E. coli* (black circle), *E. coli* (ΔLacY) (blue star), *E. coli* (ΔLacY) with SecY (ΔP) and SCVE (purple diamond), *E. coli* (ΔLacY) with SecY (ΔP, IIIGII) and SCVE (dark yellow triangle), *E. coli* (ΔLacY) with SecY (ΔP, IIIGIG) and SCVE (red square). Error bars, s.d., n = 3. **B.** The down-view 3D structures of SecY (ΔP), SecY (ΔP, IIIGII), and SecY (ΔP, IIIGIG). PDBID 3J46 was used as a template in homology modeling (Modeller v9.18). The pore-ring residues of SecY (ΔP): Ile 82, Ile 86, Ile 187, Ile 191, Ile 278, Ile 408. SecY (ΔP, IIIGII): Ile 82, Ile 86, Ile 187, Gly 191, Ile 278, Ile 408. SecY (ΔP, IIIGIG): Ile 82, Ile 86, Ile 187, Gly 191, Ile 278, Gly 408. **C.** 2-NBDG transport by *E. coli* (ΔPtsG) with SCVE and different SecY channels. The 2-NBDG uptake rate by *E. coli* (ΔPtsG) was used as a background, since 2-NBDG can also be imported via the glucose transport systems. Error bars, s.d., n = 3.

### The benefits of using the SecY channel mutant for sugar transport

Unlike free diffusion which is linear in the concentration difference, the rate of active transport or facilitated diffusion tends to approach saturation as the external sugar content increases, thus exhibiting non-linear kinetics and limiting the cell growth rate. More seriously, an excessively high sugar gradient across the cell membrane may force the cell to lose water, causing significant growth inhibition. As shown in Fig 5A-C, the specific growth rates of *E. coli* at 50, 100, and 150 g/L of glucose were 0.137, 0.127, and 0.026 h^−^ 1, while the rates of the *E. coli* (ΔPtsG) with SecY (ΔP), SCVE and Glk were 0.175, 0.228, and 0.184 h^−1^, respectively. It indicates that use of the SecY (ΔP) channel was able to improve the cell growth at high sugar levels. Similarly, *Clostridium acetobutylicum*, a Gram-positive, anaerobic bacterium, was found to grow faster when the SecY (ΔP) channel was employed for xylose transport rather than using the native XylT at 70 g/L of xylose (Fig 5D). The specific growth rate increased by over 18%, from 0.364 h^−1^ to 0.431 h^−1^. Moreover, Fig 5E and F suggest that the SecY (ΔP) channel was able to reduce the competitive inhibition caused by the glucose analogs, methyl-α-D-glucoside (α-MG) and 2-deoxy-D-glucose (2-DG), on transport of glucose in *E. coli*. Thereafter, simultaneous utilization of mixed monosaccharides by the channel-engineered *E. coli* was tested. As expected, the wild-type *E. coli* exhibited CCR when grown in the presence of glucose and xylose (Fig 5G). Intensive efforts have revealed that the PTS of the preferred carbon source plays a major role in CCR regulations (Deutscher *et al*, 2006; Görke & Stülke, 2008). However, deletion of the glucose PTS may lead to a reduced glucose uptake (Kim *et al*, 2015) (Fig 5H). Our results show that replacing the glucose PTS with SecY (ΔP), SCVE and Glk in *E. coli* enabled the cells to simultaneously utilize glucose and xylose (Fig 5I), overcoming the CCR phenomenon, and the consumption rates of both sugars were accelerated as compared to the non-channel-engineered *E. coli* (ΔPtsG), increasing the specific growth rate from 0.174 h^−1^ to 0.309 h^−1^.

**Figure 5.**
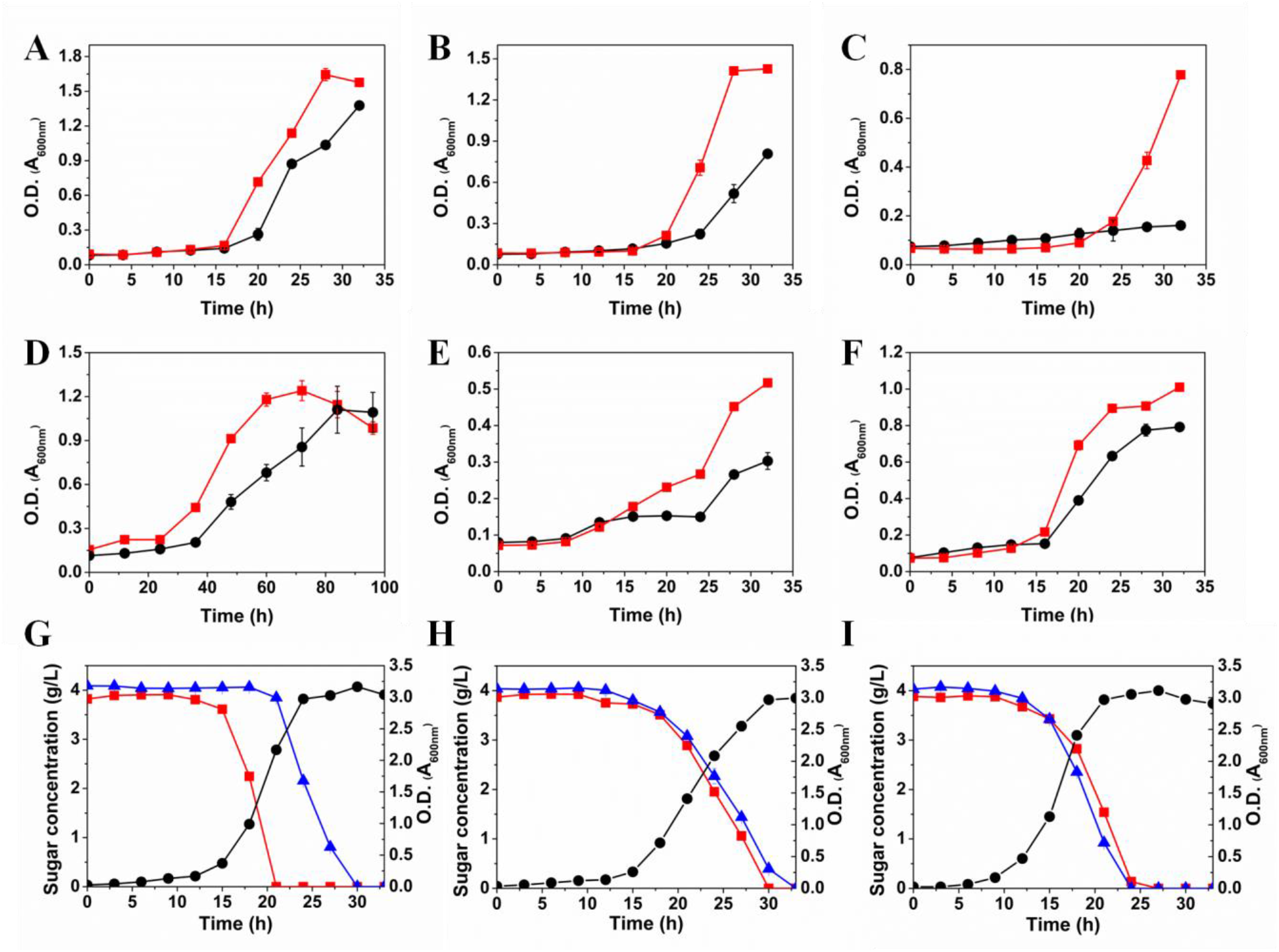
The benefits of using SecY(ΔP) for sugar transport. **A, B, C, E, F.** M9 medium, wild-type *E. coli* (black circle), *E. coli* (ΔPtsG) with SecY (ΔP), SCVE and Glk (red square). Sugars: (A) 50 g/L glucose; (B) 100 g/L glucose; (C) 150 g/L glucose; (E) 4 g/L glucose and 4 g/L 2-DG; (F) 4 g/L glucose and 12.5 g/L α-MG. Error bars, s.d., n = 3. **D.** P2 medium, 70 g/L xylose, wild-type *C. acetobutylicum* (black circle), *C. acetobutylicum* (ΔXylT) with SecY (ΔP) (red square). Error bars, s.d., n = 3. **G, H, I.** M9 medium, cell density (black circle), glucose (red square), and xylose (blue triangle). Strains: (g) wild-type *E. coli;* (h) *E. coli* (ΔPtsG); (i) *E. coli* (ΔPtsG) with SecY (ΔP), SCVE, and Glk. Error bars, s.d., n = 3.

## Materials and Methods

### Strains, vectors, and media

The strains used in this study were *E. coli* Top10 and *E. coli* BL21 (DE3), which have the genotypes: F^−^ *mcrA* Δ(*mrr-hsd*RMS-*mcr*BC) φ80 *lac*ZΔM15 Δ*lac*X74 *recA*1 *ara*Δ139 Δ(*ara-leu*)7697 *gal*U *gal*K *rps*L(Str^R^) *endA*1 *nup*G, and F^−^ *omp*T *hsd*S(r_B_^−^ m_B_^−^) *gal dcm* (DE3), respectively. *E. coli* Top10 was used only for plasmid construction. *C. acetobutylicum* ATCC 55025 was from American Type Culture Collection (USA). The λ Red recombination system (pKD13, pKD46, and pCP20) was offered by Prof. Ping-Fang Tian (Beijing University of Chemical Technology). The plasmids pETDuet-1 (ampicillin resistance), pRSFDuet-1 (kanamycin resistance), and pACYCDuet-1 (chloramphenicol resistance) were purchased from Novagen (Merck Millipore), which can be used together in *E. coli* BL21 (DE3) for co-expression of proteins under *T7* promoters. *E. coli* ER2275 (pAN1) was from Prof. Terry Papoutsakis (University of Delaware). The plasmids pIMP1-*P_thl_* and pSY6 were provided by Prof. Yang Gu and Prof. Sheng Yang from Shanghai Institutes for Biological Sciences (CAS), respectively. *P_thl_* is the *thlA* (cac2873) promoter region of *C. acetobutylicum* ATCC 824. Luria-Bertani (LB) medium contains: 10 g/L tryptone, 5 g/L yeast extract, and 10 g/L NaCl. M9 mineral medium (without sugars) contains: 7.52 g/L Na_2_HPO_4_·2H_2_O, 3 g/L KH_2_PO_4_, 0.5 g/L NaCl, 0.5 g/L NH_4_Cl, 0.25 g/L MgSO_4_·7H_2_O, 44.1 mg/L CaCl_2_·2H_2_O, 1 mg/L biotin, 1mg/L thiamin, 50 mg/L EDTA, 8.3 mg/L FeCl_3_·6H_2_O, 0.84 mg/L ZnCl_2_, 0.13 mg/L CuCl_2_·2H_2_O, 0.1 mg/L CoCl_2_·2H_2_O, 0.1 mg/L H_3_BO_3_, and 0.016 mg/L MnCl_2_·4H_2_O. Clostridial growth medium (CGM) contains: 20 g/L glucose, 2 g/L yeast extract, 4 g/L tryptone, 0.1 g/L MgSO_4_·7H_2_O, 0.5 g/L KH_2_PO_4_, 1 g/L K_2_HPO_4_, 2 g/L (NH_4_)_2_SO_4_, 2 mg/L ZnSO_4_, 0.02 g/L CoCl_2_, 0.01 g/L MnSO_4_·H_2_O, 0.015 g/L CaCl_2_·2H_2_O, 0.015 g/L FeSO_4_·7H_2_O. P2 medium contains: 70 g/L xylose, 1 g/L yeast extract, 0.5 g/L KH_2_PO_4_, 0.5 g/L K_2_HPO_4_, 2.2 g/L ammonium acetate, 1 mg/L para-amino-benzoic acid, 1 mg/L thiamin, 0.01 mg/L biotin, 0.2 g/L MgSO_4_·7H_2_O, 0.01 g/L MnSO_4_·H_2_O, 0.01 g/L FeSO_4_·7H_2_O, 0.01 g/L NaCl.

### Gene knockout

Markerless inactivation of *ptsG* (GenBank ACT42992.1), *fruA* (GenBank ACT43919.1), *manXYZ* (GenBank ACT43641.1, ACT43642.1, ACT43643.2), *xylFGH* (GenBank ACT45218.1, CP001509.3, ACT45219.1), *araFGH* (GenBank ACT43723.1, ACT43722.1, ACT43721.2), *araE* (GenBank ACT44505.1), and *lacY* (GenBank ACT42196.1) in *E. coli* BL21 (DE3) was performed with the *λ* Red recombination system (Datsenko & Wanner, 2000). Briefly, PCR product was generated by using primers (Appendix Table S2) that are homologous to regions adjacent to the gene to be inactivated and template pKD13 carrying kanamycin resistance gene that is flanked by FLP recognition target sites. The *E. coli* BL21 (DE3) harboring pKD46 was grown in LB containing 0.1 g/L ampicillin and 4 g/L arabinose at 37 °C for 1 h. After that, the PCR product was transformed (MicroPulser, Bio-Rad, USA) into cells and made disruption of chromosomal gene on LB agar plate containing 0.1 g/L kanamycin at 37 °C. The kanamycin resistance gene in the gene-knockout mutant was eliminated by using pCP20, which encodes the FLP recombinase at 30 °C and is lost at 42 °C. Disruption of *xylT* (GenBank AAK79313.1) in *C. acetobutylicum* ATCC 55025 was performed with a group II intron-based targetron technology as described previously (Shao *et al*, 2007). Briefly, the 350 bp targetron fragment for xylT gene was amplified by using the primers shown in Appendix Table S2 and pSY6 as a template. The PCR product was then digested and ligated to pSY6, yielding pSY6-xylT which was methylated in *E. coli* ER2275 (pAN1) (Mermelstein & Papoutsakis, 1993) and electroporated into *C. acetobutylicum* ATCC 55025. The strain *C. acetobutylicum* (ΔXylT) was selected on CGM plates supplemented with 50 μg/mL erythromycin at 37 °C, and the erythromycin resistance was then lost on CGM plates without any antibiotics.

### Cloning, mutation, and plasmid construction

*sCVE* (GenBank AAP73416.1) was codon optimized and synthesized by Inovogen Tech. Co. (Beijing, China). The sequence is listed below (5’→3’): ATGTACTCGTTCGTATCTGAAGAAACCGGTACTCTGATCGTGAATTCCGTGCTGCTGTTCCTGGCGTTCGTAGTCTTCCTGCTGGTCACTCTGGCTATTCTGACCGCGCTGCGTCTGTGCGCATACTGTTGTAACATCGTAAACGTTTCCCTGGTTAAACCGACGGTATACGTATACTCTCGCGTCAAAAACCTGAACAGCTCCGAAGGTGTCCCGGACCTGCTGGTT The genes, recombinant plasmids, recombinant strains, and primers used for gene cloning and gene mutation are summarized in Appendix Tables S3-6.

### Localization analysis of SecY (ΔP) and SCVE

*E. coli* was inoculated in LB containing 100 μg/mL ampicillin and 100 μg/mL kanamycin. After cells were grown to an A_600 nm_ of 0.5 (EU-2600 UV-visible spectrophotometer, Onlab, China) at 37 °C, protein expression was induced with 0.4 mM Isopropyl β-D-1-thiogalactopyranoside (IPTG) at 25 °C for 12 h. The induced *E. coli* cells were harvested by centrifugation at 5,000 × g and washed two times with phosphate-buffered saline (PBS, pH 7.4). Cells were then resuspended in PBS at 15:1 and disrupted by sonication (JY92-IIN ultrasonic homogenizer, Scientz, China) on ice. Cellular debris was removed by centrifugation for 10 min at 11,000 × g, and the cell-free extract was then centrifuged at 210,000 × g for 1 h using Optima L-80 XP Ultracentrifuge (Beckman, USA). The pellet was resuspended in PBS containing 0.01 mM MgCl_2_ and 2 % (v/v) Triton X-100 and incubated at room temperature for 30 min to solubilize the plasma membrane, and the outer-membrane fraction was then repelleted by ultracentrifugation (Fan *et al*, 2011). The fluorescence of the obtained plasma/outer-membrane fraction was measured with F-320 fluorescence spectrometer (Gangdong, China) at *λ*_Ex_ = 395 nm.

### [^14^C]-labelled glucose transport assay

The [^14^C]-labelled glucose (5.5 mCi/mmol) was purchased from American Radiolabeled Chemicals, Inc. (USA). *E. coli* strains were induced with 0.4 mM IPTG at 25 °C for 12 h in LB supplemented with 100 μg/mL ampicillin and 100 μg/mL kanamycin. Cells were harvested by centrifugation at 5,000 × g and washed twice with PBS. After that, strains were resuspended in the same buffer at an A_600 nm_ of 1 (EU-2600 UV-visible spectrophotometer, Onlab, China). The reaction was started by addition of 60 μL of [^14^C]-labelled glucose (1 mM) into 540 μL of cell suspension at room temperature. Samples were taken at 5 min and centrifuged at 5,000 × g to remove cells. 500 μL of supernant was then mixed with 5 mL scintillation cocktail. Radioactivity was measured by using Hidex 300 SL liquid scintillation counter (Hidex, Finland).

### Analysis of promoter *P_BAD_* activation

The expression kinetics of GFP controlled by *P_BAD_* promoter on arabinose was analyzed. Briefly, the *E. coli* strains with the GFP expression cassette were pre-cultured in LB containing 100 μg/mL ampicillin, 100 μg/mL kanamycin, and 30 μg/mL chloramphenicol at 37 °C over night. Cells were then sub-inoculated into M9 medium supplemented with the same antibiotics, 0.4 mM IPTG, and 4 g/L arabinose. During the growth at 37 °C, cells were harvested by centrifugation at 5,000 × g at different time, and washed two times with PBS. They were then resuspended in PBS, and the fluorescence intensity of the cells was measured by using FACSAria II (Becton Dickinson, USA).

### 2-NBDG transport assay

2-NBDG was purchased from Sigma-Aldrich Co. LLC. (USA). *E. coli* strains were inoculated in LB containing 100 μg/mL ampicillin and 100 μg/mL kanamycin, and were induced with 0.4 mM IPTG at 25 °C for 12 h. After harvested by centrifugation at 5,000 × g and washed twice with PBS, cells were resuspended in PBS. Reaction was started by addition of 2-NBDG to 10 μM at room temperature, and samples were taken at 5 min and centrifuged at 5,000 × g to remove cells. The fluorescence of supernant was measured by using F-320 fluorescence spectrometer (Gangdong, China) at *λ*_Ex_ = 475 nm and *λ*_Em_ = 550 nm (Yoshioka *et al*, 1996).

### Fermentation

*E. coli* strains were pre-cultured in LB containing 100 μg/mL ampicillin and 100 μg/mL kanamycin at 37 °C over night, and then cells were sub-inoculated into M9 medium supplemented with the same antibiotics, 0.4 mM IPTG, and sole sugar as carbon source. In mixed-sugar fermentation, the carbon sources were glucose and xylose. The cell density (A_600 nm_) in sole-sugar fermentation was determined by using Multiskan Spectrum (Thermo Scientific, USA), while in mixed-sugar fermentation, it was analyzed via EU-2600 UV-visible spectrophotometer (Onlab, China). Sugars were measured by Dionex U300 HPLC (Dionex, USA) equipped with Aminex HPX-87H Column (Bio-Rad, USA), for which 5 mmol/L H_2_SO_4_ was used as the mobile phase. For *C. acetobutylicum*, strains were anaerobically pre-cultured in CGM supplemented with 50 μg/mL erythromycin at 37 °C for 16 h in serum bottles, and then cells were sub-inoculated into P2 medium with the same conditions. The cell density of *C. acetobutylicum* was determined by using EU-2600 UV-visible spectrophotometer (Onlab, China).

### Author contributions

L-HF designed the research work. L-HF, SM, CX (Beijing University of Chemical Technology), HM, CX (Dalian University of Technology), QG, G-QD, G-BL, C-XL, Y-NQ, M-HX, and YJ performed the experiments. L-HF, SM, CX (Beijing University of Chemical Technology), S-TY, and T-WT analyzed data. L-HF and S-TY wrote the paper. SM and CX (Beijing University of Chemical Technology) contributed equally to this work.

### Conflict of interest

The authors declare that they have no conflict of interest.

## Acknowledgments

We thank Jun-Cheng Liang and Xiang-Ping Qiu from National Institute of Metrology for help with radioisotope analysis; and Miao-Miao Wang from Tsinghua University for help with ultracentrifugation. This work was supported by funding from the National Key Research and Development of China (Grant 2016YFA0204300) and the National Natural Science Foundation of China (Grant 21776010).

